# A Multimodal Neural Network Model for Early Recurrence Prediction in Lung Adenocarcinoma

**DOI:** 10.64898/2026.05.14.725244

**Authors:** Jessica A. Patricoski-Chavez, Karma Hayek, Ritambhara Singh, Christopher G. Azzoli, Jeremy L. Warner, Ece D. Gamsiz Uzun

**Affiliations:** Center for Computational Molecular Biology, Brown University, Providence, RI; Center for Clinical Cancer Informatics and Data Science (CCIDS), Brown University, Providence, RI; Department of Pathology and Laboratory Medicine, Brown University Health, Providence, RI; Graduate Program in Biotechnology, Brown University, Providence, RI; Department of Computer Science, Brown University, Providence, RI; Brown University Health Cancer Institute, Rhode Island Hospital, Providence, RI; Department of Medicine, Division of Hematology/Oncology, Brown University, Providence, RI; Department of Biostatistics, Brown University, Providence, RI; Department of Pathology and Laboratory Medicine, Warren Alpert Medical School of Brown University, Providence, RI

## Abstract

Lung adenocarcinoma (LUAD), a subtype of non-small cell lung cancer (NSCLC), is the most common primary lung cancer worldwide. Despite advancements in early detection and treatment, up to 39% of patients develop recurrent tumors following complete resection. Currently, no widely available models exist for reliably predicting early recurrence of LUAD, which is a significant prognostic factor of post-recurrence survival. Models leveraging deep learning (DL) techniques have demonstrated notable utility in cancer recurrence prediction, particularly when used in combination with both clinical and genomic data. We developed a DL-based model, **P**redicting **L**ung **A**denocarcinoma recurrence via **S**elective **M**ultimodal **A**ttention (**PLASMA**), to predict early recurrence using clinical, mRNA expression, and mutation data from patients with primary stage I-III LUAD. Trained on The Cancer Genome Atlas (TCGA) dataset, PLASMA outperformed traditional machine learning models in predicting early recurrence in both the TCGA test set and an external validation set (TRACERx Lung), achieving area under the receiver operating characteristic curve (AUROC) scores of 85.0% and 76.5%, respectively. Our results support the potential of multimodal DL for early LUAD recurrence prediction and risk stratification.

## Introduction

Advancements in early detection and treatment of lung cancer have led to a steady decline in incidence and mortality over the past decade; however, lung cancer remains the leading cause of cancer death in the United States, responsible for more deaths than second-ranking colorectal cancer and third-ranking pancreatic cancer combined.^1^ The American Cancer Society estimates that in 2026, there will be an annual incidence of 229,410 new lung cancer cases and 124,990 lung cancer-associated deaths in the United States.^1^ Lung adenocarcinoma (LUAD) is the most prevalent histologic subtype of non-small cell lung cancer (NSCLC) and the most common primary lung cancer globally, accounting for 57% and 38% of lung cancers in women and men, respectively.^2,3^ While most NSCLC is diagnosed in the advanced or metastatic setting, increasing use of targeted NSCLC screening and other diagnostic strategies are leading to increasing rates of earlier stage, potentially curable disease. Despite advances in the diagnosis and treatment of LUAD, overall recurrence rates in non-metastatic disease remain high, with 18%-39% of patients experiencing recurrence following complete resection. Furthermore, recurrence rates vary significantly by stage, with recurrence rates for stages IA and IIIB as varied as 12% and 93%, respectively.^4–8^ Prognosis following recurrence is poor; in a cohort of patients with recurrence following complete resection of stage IA-IIIB LUAD, 2- and 5-year post-recurrence survival were 65.2% and 29.8%, respectively, underscoring the clinical need for more precise risk stratification following primary tumor treatment.^9^ In particular, personalized risk characterization of early LUAD recurrence can profoundly improve patient outcomes by facilitating timely intervention, minimizing risk of overtreatment, and optimizing treatment selection.^10,11^

Traditional methods for characterizing risk, such as the TNM staging system, rely heavily on clinicopathological factors. While TNM stage is an established correlate of prognosis, it only partially captures the biological heterogeneity of LUAD, leading to wide variations in recurrence incidence and timing within the same stage groupings.^12,13^ As such, there is a critical need for more robust predictive tools that can augment conventional risk assessment frameworks. In response to these gaps, various machine learning (ML) and deep learning (DL) models have been developed. Jones et al.^14^ integrated clinicopathologic features with summary-level genomic features and mutation data from 50 genes in order to predict recurrence free survival (RFS) in early-stage LUAD patients. Their RFS prediction model, PRecur, used ensemble gradient boosting survival trees and outperformed TNM-based models in identifying high-risk patients (concordance probability estimate, 0.73 vs 0.61). Xu et al.^15^ developed a graph neural network, IBPGNET, with embedded pathway hierarchy relationships, copy number variant data, and somatic mutation data to predict LUAD recurrence. Their model achieved an area under the precision-recall curve (AUPRC) of 0.79 and an area under the receiver operating characteristic curve (AUROC) of 0.88. Pu et al.^16^ utilized a multivariate Cox proportional hazards regression model and various ML models to predict recurrence post-surgical resection of NSCLC using clinicopathologic features and preoperative chest CT scans; their models achieved AUROCs ranging from 0.75 to 0.77. Xu et al.^17^ identified differentially expressed genes between primary and recurrent LUAD tumors and utilized univariate Cox regression to identify RFS-related gene candidates before splitting the data and applying LASSO Cox regression to determine which candidates had high RFS prediction power. Thirteen genes were manually selected from this subset for their multivariate Cox regression model. Authors reported an AUROC of 0.96 identifying patients with a high risk of LUAD recurrence; however, it is unclear if this performance is based on the training set, test set, or external validation set.

While existing LUAD recurrence models are valuable, there is a current lack of models that predict *early* LUAD recurrence, which studies have shown is a significant prognostic factor of post-recurrence survival.^9^ Additionally, we have previously demonstrated that it is possible to predict early cancer recurrence in glioma patients using clinical and genomic data from primary tumors only.^18^ Given this unmet need, we developed a DL-based early recurrence prediction model, **P**redicting **L**ung **A**denocarcinoma recurrence via **S**elective **M**ultimodal **A**ttention (**PLASMA**), using clinical, mRNA expression, and somatic mutation data from patients with primary LUAD tumor samples from The Cancer Genome Atlas (TCGA).^19,20^ Our results underscore the potential for incorporating multimodal DL classifiers with traditional approaches to early LUAD recurrence prediction.

## Methods

In this study, we utilized two publicly available LUAD datasets containing clinical and genomic information. The Cancer Genome Atlas (TCGA) is a landmark initiative systematically characterizing the clinical and molecular landscapes of 33 cancer types through the large-scale genomic profiling of more than 11,000 tumor samples.^19^ To create our first dataset, we downloaded clinical data, somatic mutations, and mRNA expression data for the PanCancer TCGA-LUAD dataset from cBioPortal (cbioportal.org).^21,22^ To minimize missingness among clinical features, we also retrieved clinical data from the University of California, Santa Cruz Xena Browser (xenabrowser.net),^23–26^ the TCGA Clinical Data Resource (CDR) outcome and follow-up files,^27^ and the Firehose Legacy release of TCGA-LUAD.^28^ As an independent external validation set, we downloaded LUAD data from TRACERx (TRAcking non-small cell lung Cancer Evolution through therapy [Rx]) Lung.^29–31^ TRACERx is a collection of multicenter, prospective longitudinal studies for patients with NSCLC (TRACERx Lung) and renal cancer (TRACERx Renal) created to investigate genomic evolution in cancer and the relationship between intratumor heterogeneity and patient outcomes.^32–34^

The TCGA and TRACERx datasets were limited to patients meeting the inclusion criteria outlined in Figure 1. For the TCGA dataset, we removed patients missing age or smoking status, patients with a censored disease-free interval (DFI) event (from the standardized CDR), patients without an explicit recurrence indicator and associated time metric, patients with stage IIIB or IV tumors, and patients missing all three data types (clinical, expression, and mutation). Explicit recurrence indicators included (i) the presence of a recurrent tumor sample in the raw dataset, (ii) a disease-free interval (DFI) event coded as 1, (iii) a new tumor event (NTE) type labeled as *Locoregional recurrence* or *Distant Metastasis*, (iv) diagnosis timeline entries with a status of *Locoregional recurrence* or *Distant Metastasis*, or (v) treatment timeline entries specifying a treatment site of *Local Recurrence* or *Distant Recurrence*, or a regimen indication of *Recurrence*. For the TRACERx dataset, we removed patients with missing recurrence time values, patients with stage IIIB or IV tumors, and patients with tumors for which no tumor regions passed the sequencing quality-control checks outlined by Martínez-Ruiz et al.^29,31^ (Figure 1). For TRACERx, time to recurrence (TTR) was pre-calculated and provided by a single variable (Recurrence_time_use). For TCGA patients, TTR was derived by prioritizing available TTR-related variables across three tiers: (1) start dates of distant recurrence treatment, local recurrence treatment, recurrence regimen indication, and locoregional recurrence diagnosis; (2) start date of distant metastasis diagnosis; and (3) days to NTE and DFI time. For each patient, TTR was defined as the earliest non-null value within the highest-priority non-empty tier. In both datasets, patients with TTR values below the TCGA median (434 days) were classified as *early* recurrence, and those with values greater than or equal to the median as *late* recurrence.

**Figure 1.**
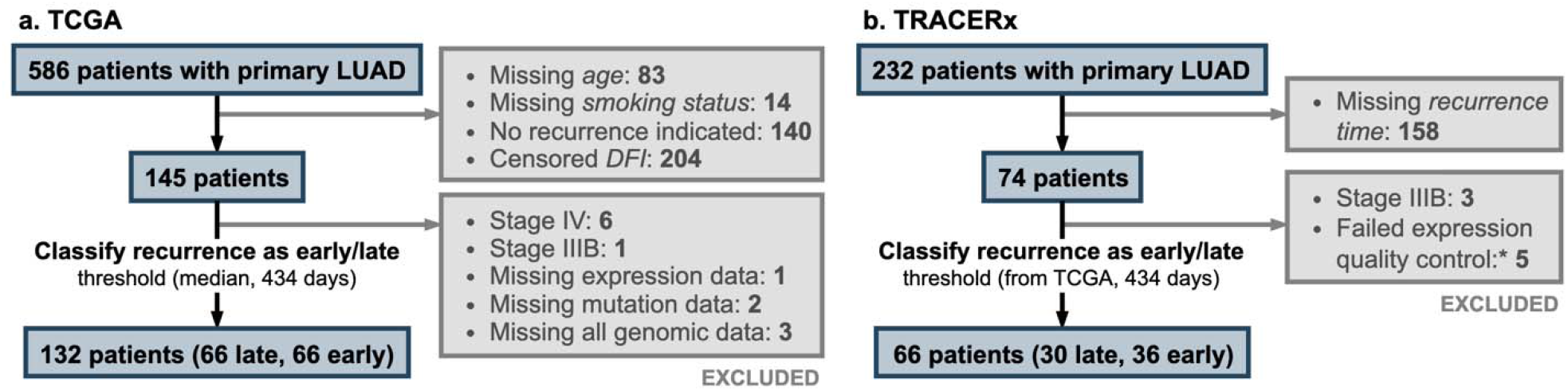
Patient Cohorts. Inclusion and exclusion criteria for the (**a**) TCGA and (**b**) TRACERx cohorts.

### Data Preprocessing

After applying the inclusion criteria in Figure 1, genomic data features (genes) were limited to currently approved protein-coding genes on chromosomes 1-22, according to the HUGO Gene Nomenclature Committee BioMart Gene repository.^35^To minimize exclusion of qualifying genes, gene features were mapped to their approved symbol using the provided gene symbols, alias symbols, and former symbols. TRACERx gene expression data (RSEM counts) were upper-quartile (75th percentile) scaled per sample to 1,000 to match the normalization strategy already applied to the TCGA data. Gene expression data for both cohorts were then log_2_(*x*+1) transformed, where *x* represents the normalized RSEM counts. Using mutation annotation data, mutation features were created by calculating the number of non-silent mutations per gene per patient. For the distribution of variant classification types, see Supplementary Methods S1 and Supplementary Table S1. However, analyses of the TCGA training set mutation features revealed 99.98% of genes had a median mutation count of zero, and only 2.84% of genes had a maximum count ≥3. Given this substantial skew, mutation counts were binarized to represent the presence or absence of one or more mutations in a given gene. The resulting genomic features for each patient were mRNA expression per gene and binarized mutation counts per gene. All genomic and clinical features were then limited to those present in both datasets. The resulting feature space included mRNA expression per gene, binarized mutation counts per gene, age at diagnosis, initial pathologic stage (categorically encoded stages I-III), sex, current smoking status (0: non-smoker, 1: smoker), and smoking history (0: never-smoker, 1: ever-smoker). The TCGA cohort was partitioned into training, validation, and testing using a 70/15/15 split stratified by outcome label (Figure 2).

**Figure 2.**
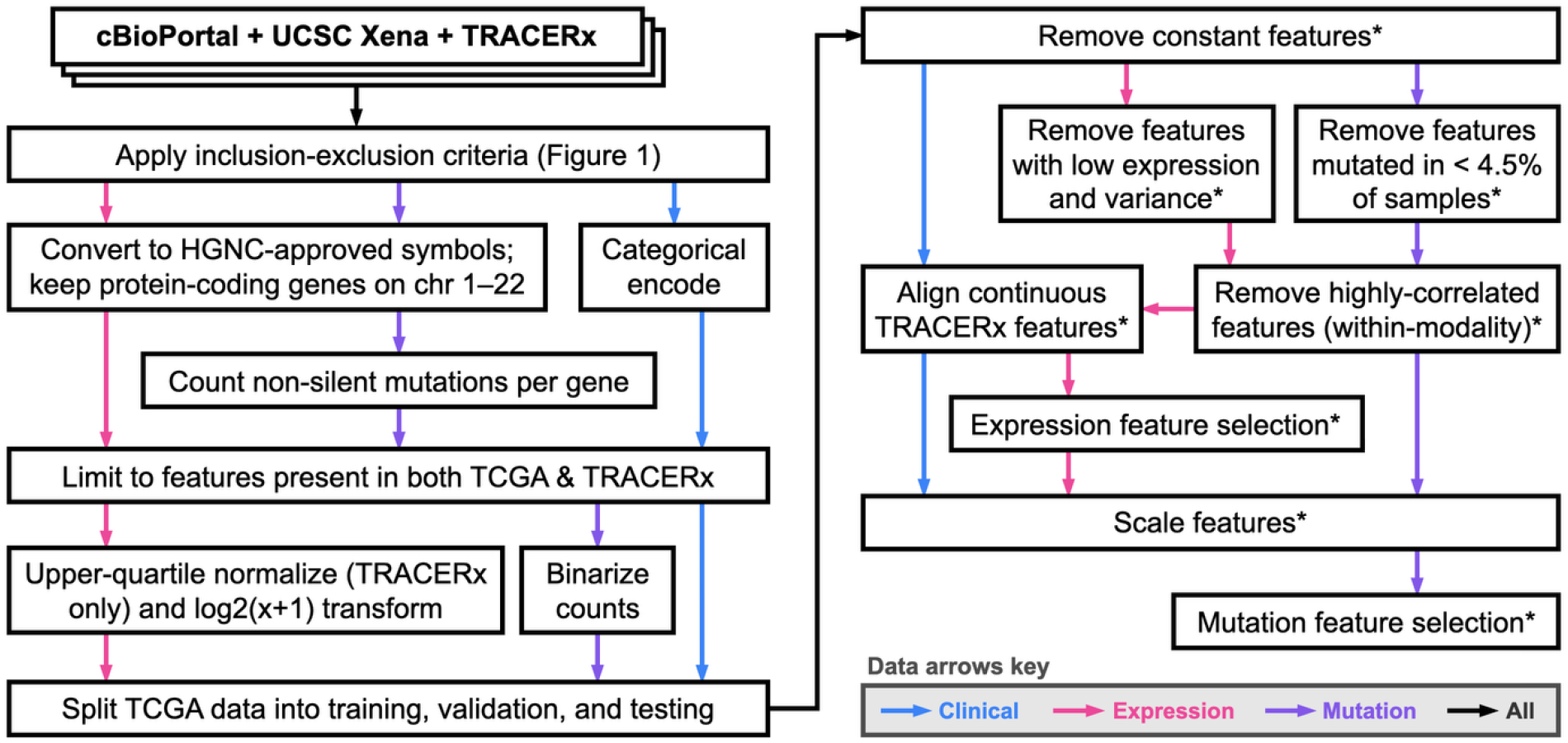
Data preprocessing pipeline. *Preprocessing steps were fit on the TCGA training set only.

Fit on the TCGA training set, a filtering pipeline removed (i) constant features, (ii) mutation features with nonsynonymous variants in <4.5% of samples, (iii) expression features with low expression values (>1 count in <20% of samples or variance <0.01), (iv) expression features with low variance (25th percentile), and (v) within-modality redundancies in the form of highly-correlated features (correlation ≥0.95). For further details on the correlation analysis, see Supplementary Methods S2. A threshold of 4.5% was chosen based on the distribution of mutation frequencies in the training set with the intention of minimizing sparsity and noise (Supplementary Figure S1). This pipeline resulted in 7 clinical features (unchanged), 11,686 expression features, and 1009 mutation features. Next, expression features were filtered in an unsupervised manner to ensure comparability between the primary (TCGA) and validation (TRACERx) cohorts without data leakage. Expression features were excluded if they exhibited near-zero variance in the target dataset (<0.001) or if their mean expression differed by more than 3.5 standard deviations between datasets, suggesting potential batch effects or systematic distributional discordance that could confound generalization. Expression feature selection consisted of a signal-to-noise ratio-based univariate feature selector, incorporating stability selection (75 stratified bootstraps at 85% subsample) and a literature-derived gene prior via rank-based bonus (for the complete list of literature-derived genes, see the Supplementary Methods S3); the top 50 expression features were retained. Prior to scaling, continuous features (expression and age at diagnosis) in TRACERx were aligned to the TCGA train distribution using unsupervised quantile domain alignment. Next, scalers were selected and applied per-feature for all continuous features according to outlier fraction and skew. Lastly, 50 mutation features were selected using an eXtreme Gradient Boosting (XGBoost) classifier^36^ with a gene prior rank-based bonus (Figure 2). Selecting the top 50 features for each genomic modality type provided a balanced number of LUAD-relevant genes while maintaining a manageable feature space and mitigating the risk of overfitting.

### Final Cohorts

The TCGA dataset contained 132 patients meeting all criteria outlined in Figure 1 (Table 1). Of those 132 patients, 86.4% (*n* = 114) were current or former smokers. Patients were most commonly diagnosed with stage I tumors (IA: *n* = 29 [22.0%], IB: *n* = 37 [28.0%]), followed by stage II (IIA: *n* = 14 [10.6%], IIB: *n* = 30 [22.7%]) and stage III (IIIA: *n* = 22 [16.7%]). Median age at diagnosis was 66.0 years, and 58.3% of patients were female (*n* = 77). The TRACERx dataset contained 66 patients meeting all criteria. Of those 67 patients, 54.5% (*n* = 36) had an early recurrence, 45.5% (*n* = 30) had a late recurrence, and 93.9% (*n* = 62) were current or former smokers. Unlike TCGA, patients were most commonly diagnosed with stage III tumors (IIIA: *n* = 25 [37.9%]), followed by stage I (IA: *n* = 5 [7.6%], IB: *n* = 17 [25.8%]) and stage II (IIA: *n* = 11 [16.7%], IIB: *n* = 8 [12.1%]). Median age at diagnosis was 70.0 years, and 47.0% of patients were female (*n* = 31). Descriptive and demographic statistics for the TCGA and TRACERx cohorts are available in Tables 1 and 2, respectively.

**Table 1.** Descriptive statistics for the TCGA and TRACERx patient cohorts, according to recurrence outcome.

|  |  | TCGA |  |  | TRACERx |  |  |
| --- | --- | --- | --- | --- | --- | --- | --- |
|  |  | All (132) | Early (66) | Late (66) | All (66) | Early (36) | Late (30) |
| Sex [N (%)] | Female | 77 (58) | 35 (53) | 42 (64) | 31 (47) | 12 (33) | 19 (63) |
|  | Male | 55 (42) | 31 (47) | 24 (36) | 35 (53) | 24 (67) | 11 (37) |
| **Stage [N (%)]** | IA | 29 (22) | 10 (15) | 19 (29) | 5 (8) | 1 (3) | 4 (13) |
|  | IB | 37 (28) | 14 (21) | 23 (35) | 17 (26) | 5 (14) | 12 (40) |
|  | IIA | 14 (11) | 8 (12) | 6 (9) | 11 (17) | 8 (22) | 3 (10) |
|  | IIB | 30 (23) | 17 (26) | 13 (20) | 8 (12) | 6 (17) | 2 (7) |
|  | IIIA | 22 (17) | 17 (26) | 5 (8) | 25 (38) | 16 (44) | 9 (30) |
| **Smoking status [N (%)]** | Former | 88 (67) | 50 (76) | 38 (58) | 41 (62) | 24 (67) | 17 (57) |
|  | Current | 26 (20) | 9 (14) | 17 (26) | 21 (32) | 11 (31) | 10 (33) |
|  | Never | 18 (14) | 7 (11) | 11 (17) | 4 (6) | 1 (3) | 3 (10) |
| **Age [yr]** | Median | 66.0 | 66.0 | 67.0 | 70.0 | 71.0 | 68.0 |
|  | IQR | 60.0-72.0 | 59.0-72.0 | 61.0-73.0 | 64.0-76.0 | 64.0-77.3 | 64.0-73.0 |
| **TTR [days]** | Median | 434.0 | 244.0 | 659.0 | 369.0 | 210.5 | 682.0 |
|  | IQR | 245.0-656.5 | 183.3-326.0 | 502.3-878.5 | 206.8-655.0 | 167.3-307.8 | 537.8-930.5 |

**Table 2.** Demographic data for the TCGA and TRACERx patient cohorts, according to recurrence outcome.

| TCGA* |  | All (132) | Early (66) | Late (66) | TRACERx |  | All (66) | Early (36) | Late (30) |
| --- | --- | --- | --- | --- | --- | --- | --- | --- | --- |
| **Race [N (%)]** | American Indian or Alaska Native | 1 (<1) | 1 (2) | 0 (0) | **Ethnicity [N (%)]** | White - British | 57 (86) | 27 (73) | 30 (100) |
|  | Asian | 2 (2) | 2 (3) | 0 (0) |  | White - European | 3 (5) | 3 (8) | 0 (0) |
|  | Black or African American | 14 (11) | 5 (8) | 9 (14) |  | White - Irish | 2 (3) | 2 (6) | 0 (0) |
|  | White | 105 (80) | 53 (80) | 52 (79) |  | White - Other | 1 (2) | 1 (3) | 0 (0) |
| **Ethnicity [N (%)]** | Hispanic or Latino | 1 (<1) | 0 (0) | 1 (2) |  | Caribbean | 1 (2) | 1 (3) | 0 (0) |
|  | Not Hispanic or Latino | 107 (81) | 53 (80) | 54 (82) |  | Middle Eastern | 1 (2) | 1 (3) | 0 (0) |
|  |  |  |  |  |  | Indian | 1 (2) | 1 (3) | 0 (0) |

\*Race and ethnicity were unavailable for 10 and 24 TCGA patients, respectively.

### PLASMA

We developed our model, **P**redicting **L**ung **A**denocarcinoma recurrence via **S**elective **M**ultimodal **A**ttention (**PLASMA**), using clinical, mRNA expression, and mutation data from patients with primary LUAD tumor samples. The PyTorch^37^ model framework is shown in Figure 3. Each data type is first passed through its own feed-forward neural network (FFNEncoder), consisting of one or more fully-connected layers with layer normalization, a nonlinear activation function, and dropout. The role of each encoder is to compress its raw input into a compact, fixed-length numerical vector (an *embedding*) that captures the most informative aspects of that modality. We use modality-specific encoders rather than a shared encoder because the inputs differ substantially in dimensionality, scale, and biological meaning, and learning a separate transformation for each allows the network to extract patterns appropriate to its data type. Layer normalization stabilizes training across the very different input distributions, dropout reduces overfitting, and the nonlinear activation enables the encoder to learn non-trivial relationships between features.

**Figure 3.**
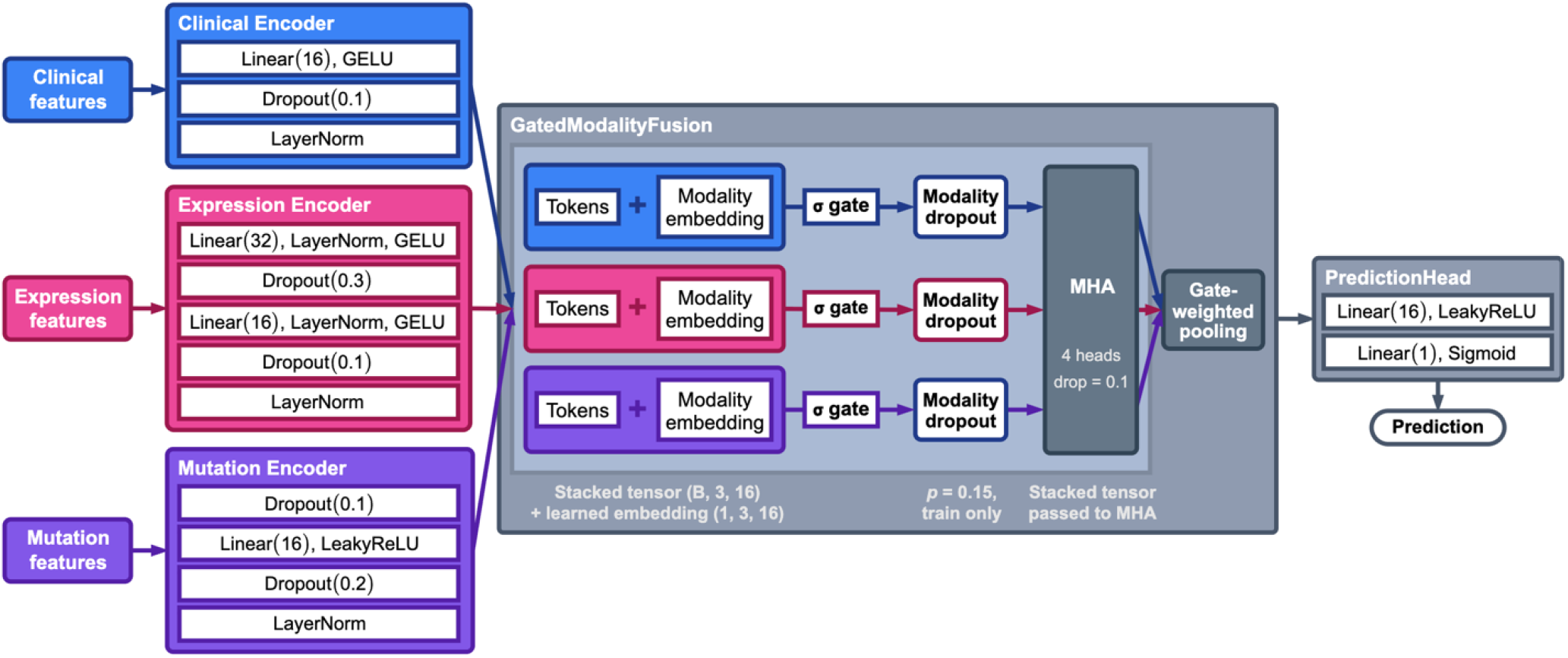
PLASMA Model Architecture.

The three resulting embeddings are then treated as tokens (individual vector elements in a short sequence) and passed to the gated multi-head self-attention fusion module (GatedModalityFusion). Because self-attention is inherently permutation-invariant, we add a learned modality-type embedding to each token so the model can distinguish clinical tokens from mRNA tokens from mutation tokens. The module then computes a sigmoid confidence gate for each modality, which allows the model to dynamically up-weight modalities carrying a strong recurrence signal and suppress those that are noisy or uninformative for that patient (hence *selective* attention). Self-attention is then applied across the gated tokens, allowing each modality’s representation to be refined based on context from the others. Finally, the three attended tokens are combined into a single fused embedding via gate-weighted pooling, so that more confident modalities contribute more strongly to the final representation. The subsequent embedding then passes through a small post-attention adapter (a linear layer with GELU activation and layer normalization) that refines the pooled representation before prediction. This fused embedding is passed to a fully-connected prediction head (PredictionHead) that outputs the recurrence prediction.

During training, we additionally applied modality dropout at a rate of 15%, wherein entire modality tokens were randomly zeroed before fusion. This forced the model to make sensible predictions even when a modality was absent, thus improving robustness to missing modalities and preventing the model from relying too heavily on any single data source. Each model component was trained with its own Adam optimizer. Learning rates were adapted using ReduceLROnPlateau schedulers, and all weights were initialized using Xavier uniform initialization to maintain stable activation variance across layers at the start of training. Models were trained for a maximum of 30 epochs with a batch size of 12, utilizing early stopping after a grace period of 12 epochs without validation improvement.

To combat our small sample size and reduce sensitivity to random initialization, the full training and evaluation procedure was repeated across five random seeds, and per-seed output logits were averaged to form an ensemble prediction. Post-hoc probability calibration was performed on the ensemble validation logits using Platt scaling (logistic regression) and applied to all held-out sets (TCGA testing and TRACERx). The classification threshold was selected once on the calibrated validation set probabilities by maximizing the geometric mean of sensitivity and specificity. Hyperparameters were selected via stratified k-fold cross-validation (CV; k=3) on a development pool comprising the merged TCGA training and validation splits, with the held-out TCGA test set and the external TRACERx cohort left unseen during tuning. Within each fold, 15% of the training split was designated as an inner validation set to provide signals for early stopping and learning-rate scheduling. Configurations were ranked by mean held-out AUROC across folds. Further details on the hyperparameter tuning are provided in Supplementary Methods S4.

### Model Evaluation

To assess the model performance, we benchmarked PLASMA against the following traditional classifiers: 1) support vector machine (SVM); 2) multi-layer perceptron (MLP); 3) logistic regression (LR); and 4) k-nearest neighbors (KNN); and 5) random forest (RF).^38^ Baseline models were evaluated using the same multiseed training (for stochastic baselines), calibration, and threshold selection used for PLASMA. For each model, we evaluated performance using AUROC, AUPRC, balanced accuracy, average precision, recall, specificity, F1-score, true negatives, false positives, false negatives, and true positives. Baseline classifiers were tuned using exhaustive grid search over their respective hyperparameter spaces, and evaluated on the same stratified 3-fold CV protocol applied to PLASMA, including identical patient cohorts, fold assignments, and inner-validation carve-out (Supplementary Methods S4). To assess the importance of each modality type on PLASMA predictive performance, we conducted an ablation analysis with PLASMA models trained on every possible combination of the three input modalities, as well as a model using simple concatenation in place of GatedModalityFusion, all tuned using the same hyperparameter strategy. To assess the importance of individual features, we utilized the PLASMA model from the first seed together with SHapley Additive exPlanations (SHAP).^39^ SHAP is a model interpretation framework that assigns each input feature a Shapley value representing its average marginal contribution to a model’s prediction. Positive SHAP values increase the model’s predicted likelihood of recurrence relative to the baseline prediction, while negative SHAP values indicate features that decrease it. In this study, we utilize SHAP PermutationExplainer, which is a model-agnostic method that estimates SHAP values by iterating through random feature permutations and measuring the change in prediction. We selected PermutationExplainer as it makes no assumptions about a model’s architecture or the differentiability of its components.

## Results

On the TCGA testing set, PLASMA outperformed all baseline models in AUROC (85.0%), AUPRC (87.9%), balanced accuracy (80.0%), average precision (88.5%), specificity (80.0%), F1 score (80.0%), true negatives (*n* = 8), and false positives (*n* = 2). PLASMA tied for best recall (80.0%), false negatives (*n* = 2), and true positives (*n* = 8) with LR, SVM, and RF. However, these baselines demonstrated notably worse performance in all other metrics, particularly specificity (LR: 40.0%, SVM: 30.0%, RF: 50.0%), balanced accuracy (LR: 60.0%, SVM: 55.0%, RF: 65.0%), and AUROC (LR: 67.0%, SVM: 65.0%, RF: 62.0%).

On the TRACERx external validation set, PLASMA outperformed all baseline models in AUROC (76.5%), AUPRC (79.6%), balanced accuracy (71.1%), average precision (80.0%), specificity (86.7%), true negatives (*n* = 26), and false positives (*n* = 4). RF achieved superior recall (77.8%), F1 score (70.0%), false negatives (*n* = 8), and true positives (*n* = 28). However, RF showed substantially worse performance in all other metrics, including specificity (46.7%), AUROC (58.8%), AUPRC (55.6%), average precision (57.0%), true negatives (*n* = 14), and false positives (*n* = 16). Full performance metrics for all models evaluated on the TCGA testing set and TRACERx external validation set are provided in Table 3. Receiver operating characteristic (ROC) curves and precision-recall (PR) curves for all models are displayed in Figure 4. Training and validation loss curves for all seeds are provided in Supplementary Figure S2.

**Table 3.** Performance metrics for PLASMA and the baseline comparison models on the TCGA testing and TRACERx datasets. The bolded metric is the best-in-class, stratified by dataset.

|  |  | AUROC | AUPRC | Acc | Precision | Recall | Specificity | F1 | TN | FP | FN | TP |
| --- | --- | --- | --- | --- | --- | --- | --- | --- | --- | --- | --- | --- |
| **TCGA** | **PLASMA** | **85.0** | **87.9** | **80.0** | **88.5** | **80.0** | **80.0** | **80.0** | **8** | **2** | **2** | **8** |
|  | **KNN** | 74.0 | 75.3 | 65.0 | 76.6 | 70.0 | 60.0 | 66.7 | 6 | 4 | 3 | 7 |
|  | **LR** | 67.0 | 71.7 | 60.0 | 73.3 | **80.0** | 40.0 | 66.7 | 4 | 6 | **2** | **8** |
|  | **MLP** | 66.0 | 76.6 | 50.0 | 77.6 | 60.0 | 40.0 | 54.6 | 4 | 6 | 4 | 6 |
|  | **SVM** | 65.0 | 73.6 | 55.0 | 74.7 | **80.0** | 30.0 | 64.0 | 3 | 7 | **2** | **8** |
|  | **RF** | 62.0 | 66.2 | 65.0 | 68.1 | **80.0** | 50.0 | 69.6 | 5 | 5 | **2** | **8** |
| **TRACERx** | **PLASMA** | **76.5** | **79.6** | **71.1** | **80.0** | 55.6 | **86.7** | 66.7 | **26** | **4** | 16 | 20 |
|  | **KNN** | 52.6 | 52.4 | 55.6 | 54.3 | 61.1 | 50.0 | 60.3 | 15 | 15 | 14 | 22 |
|  | **LR** | 53.9 | 52.3 | 59.7 | 53.7 | 69.4 | 50.0 | 65.8 | 15 | 15 | 11 | 25 |
|  | **MLP** | 61.0 | 62.0 | 57.2 | 63.0 | 61.1 | 53.3 | 61.1 | 16 | 14 | 14 | 22 |
|  | **SVM** | 61.4 | 61.6 | 62.5 | 62.8 | 75.0 | 50.0 | 69.2 | 15 | 15 | 9 | 27 |
|  | **RF** | 58.8 | 55.6 | 62.2 | 57.0 | **77.8** | 46.7 | **70.0** | 14 | 16 | **8** | **28** |

Acc = balanced accuracy; TN = true negatives; FP = false positives; FN = false negatives; TP = true positives.

**Figure 4.**
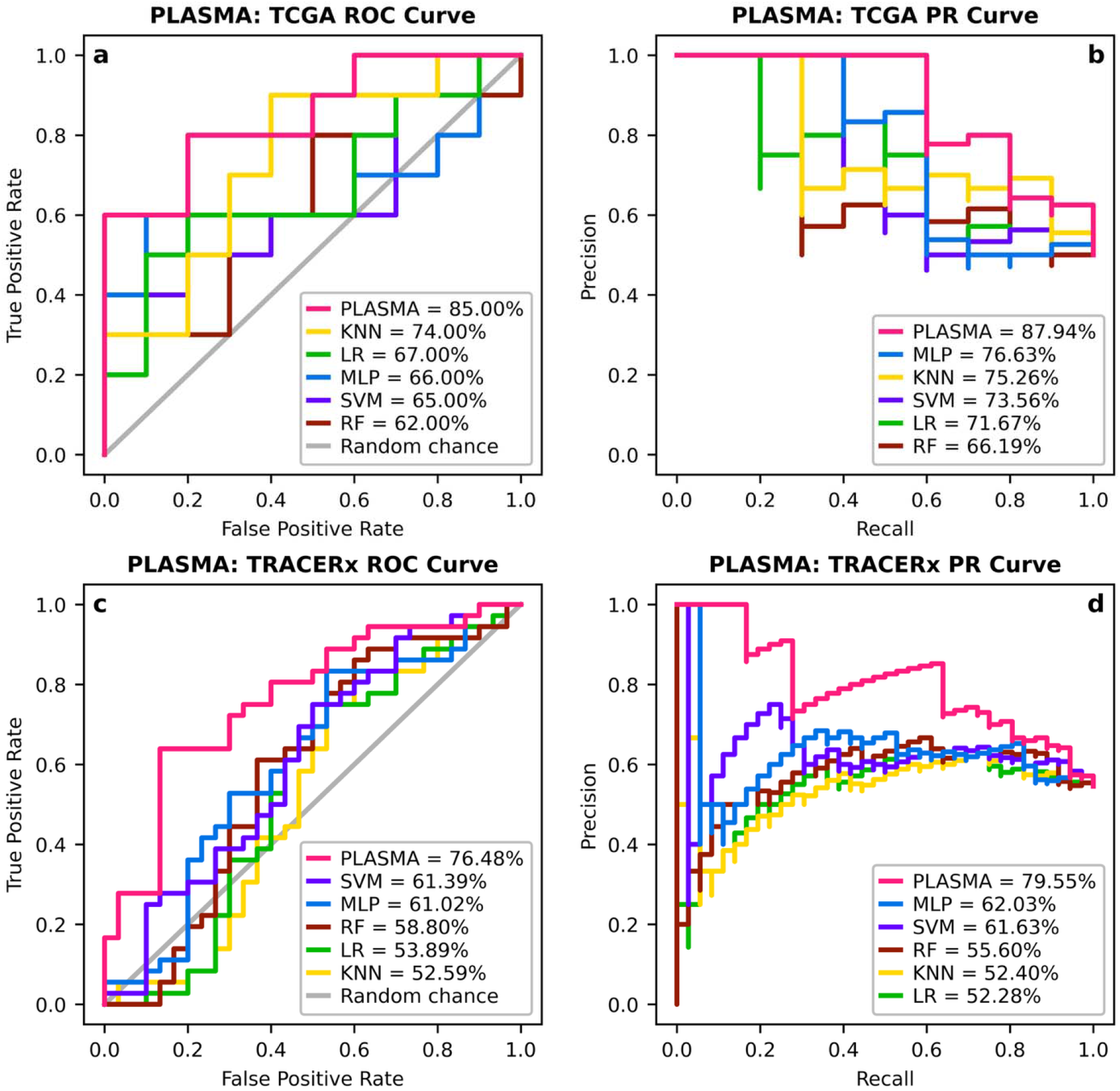
Baseline Performance Comparison. ROC and PR curves comparing PLASMA and the traditional baseline models’ performance on the TCGA test cohort (**a, b**) and TRACERx cohort (**c, d**).

The increased performance of PLASMA over the baselines may reflect a set of architectural choices well-matched to multimodal recurrence prediction. Modality-specific encoders accommodate the very different dimensionality, scale, and feature semantics of clinical, mRNA, and mutation data rather than forcing them into a shared representation prematurely. The sigmoid confidence gates perform per-sample reweighting of modalities, which we hypothesize is valuable given the well-documented molecular heterogeneity of LUAD; if the most informative modality varies across tumors, a model that can adapt its modality weighting on a per-sample basis has more flexibility than one that applies a fixed weighting learned across the cohort. Self-attention enables the representation of each modality to be conditioned on the others, while modality dropout likely acts as a useful regularizer given the low-sample, high-dimensional training regime. Collectively, these inductive biases (selective weighting, cross-modality reasoning, and robustness regularization) are largely absent from the baselines and may account for much of the observed performance gap.

To assess whether each modality ablation exceeded chance-level discrimination, we performed a one-sided label-permutation test of AUROC (1,000 permutations) on the ensemble-calibrated probabilities for each test cohort, with Bonferroni correction applied across the seven variants tested per cohort. After correction, two PLASMA models achieved significant discrimination above chance in both the TCGA test cohort (*n* = 20) and the independent TRACERx cohort (*n* = 66): the full multimodal model (TCGA: AUROC = 0.850, Bonferroni p = 0.021; TRACERx: AUROC = 0.765%, Bonferroni p < 0.001) and the clinical-only model (TCGA: AUROC = 0.820, Bonferroni p = 0.035; TRACERx: AUROC = 0.697, Bonferroni p = 0.035). Replacing GatedModalityFusion with a simple concatenation and feed-forward MLP eliminated this performance: the concatenation variant failed to exceed chance on either cohort (TCGA: AUROC = 0.650, Bonferroni p = 0.294; TRACERx: AUROC = 0.533, Bonferroni p = 0.618). Full test statistics, 95% confidence intervals, effect sizes, and per-variant p-values for all comparisons are reported in Supplementary Methods S5 and Tables S5-S6.

Two modality ablations, clinical + mutation (CM) and expression + mutation (EM), yielded AUROCs substantially below chance on the TCGA test cohort (0.36 and 0.23, respectively), whereas both variants performed considerably better on the larger TRACERx cohort, with CM clearing corrected significance (AUROC = 0.71) and EM producing a point estimate above chance though not surviving Bonferroni correction (AUROC = 0.64). This discrepancy is best attributed to small-sample (*n* = 20) variance rather than systematic anti-prediction: in a cohort of 20 patients, a small number of inverted predictions can shift AUROC by 10–20 percentage points, a pattern reflected in the wider confidence intervals observed for TCGA test estimates throughout (Supplementary Table S5). The full multimodal model, by contrast, achieved corrected significance on both cohorts (TCGA: AUROC = 0.85; TRACERx: AUROC = 0.77), indicating that fusion of all three modalities leads to greater robustness to cohort-specific patient distributions.

The decrease in performance when we replaced GatedModalityFusion with simple concatenation suggests that the fusion mechanism, rather than merely the presence of all three modalities, is critical to PLASMA’s predictive ability. We hypothesize several contributing factors. First, concatenation forces the downstream PredictionHead to learn cross-modality interactions implicitly through a single dense layer operating on a high-dimensional joint vector, which substantially increases parameter count and creates a harder optimization problem given our small sample size. As a result, the model is likely unable to identify informative cross-modality patterns before overfitting to spurious patterns. Second, the simple concatenation treats every dimension of each modality as equally relevant for each patient, with no mechanism to down-weight noisy or uninformative modalities on a per-sample basis, unlike GatedModalityFusion, which uses sigmoid confidence gates to suppress modalities that carry little signal for a given patient. Finally, concatenation provides no mechanism for modalities to contextualize one another, whereas self-attention allows the prognostic weight of a feature in one modality to be conditioned on the state of the others.

Median per-feature SHAP importance, calculated on the TCGA testing set, was highest for clinical features (0.128), followed by mutation features (0.025), and expression features (0.024). A summary plot outlining the 20 most influential features to PLASMA recurrence predictions is shown in Figure 5. The most significant feature from each modality type included pathologic tumor stage, *TP53* mutations, and *INSRR* expression.

**Figure 5.**
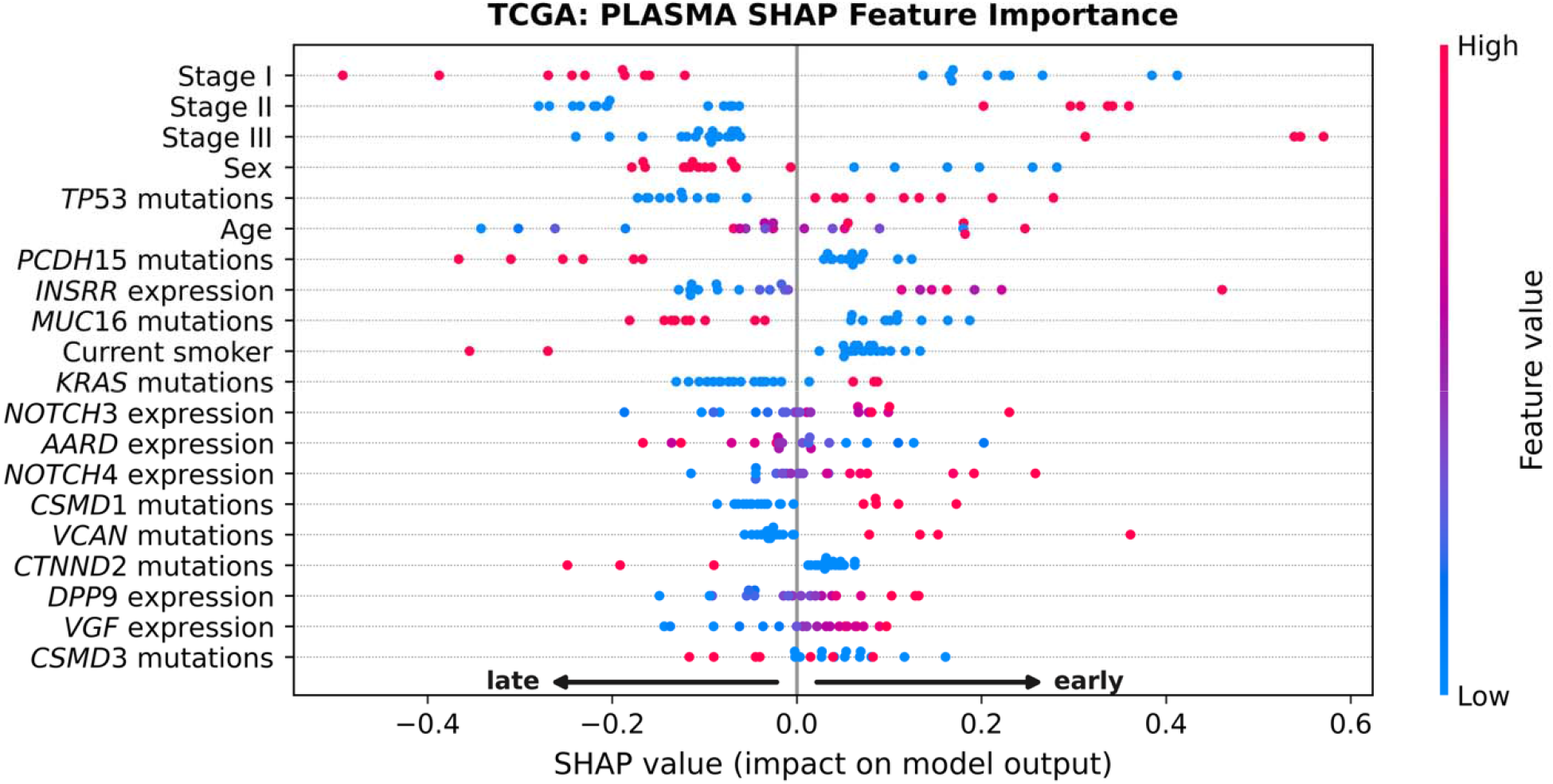
SHAP Feature Importance. The 20 most influential features to PLASMA, according to SHAP PermutationExplainer and the TCGA testing set. Positive SHAP values (dots to the right) indicate an increase in the model’s output (towards an early prediction) and negative values (dots to the left) indicate a decrease (towards a late prediction). For patient sex, red dots represent female patients.

## Discussion

PLASMA, a multimodal DL classifier, surpassed baseline comparators in predicting LUAD recurrence in both the TCGA and TRACERx datasets, across most standard metrics of performance. Multiple features of high SHAP importance have established associations with lung adenocarcinoma prognosis. For example, SHAP PermutationExplainer indicates that PLASMA was influenced towards early recurrence predictions by stage II and stage III tumors, increased patient age, male patients, *TP53* mutations, *KRAS* mutations, and high expression of *DPP9, VGF, NOTCH3*, and *NOTCH4*. Conversely, PLASMA was influenced towards late recurrence by stage I tumors.

While recurrence incidence and timing can vary within tumor stage groupings, advanced tumor stage is a well-established factor in LUAD outcomes. In a cohort of 426 patients with completely resected stage I-III LUAD treated at Memorial Sloan Kettering Cancer Center, patients with stage II and stage III tumors experienced significantly worse RFS.^40^A systematic literature review and meta-analysis of 85 recurrence studies enrolling patients with stage I-III NSCLC found that advanced tumor stage (stages II and III), older age, and a higher percentage of male patients were significantly associated with worse RFS across time points.^41^ *TP53*, one of the most commonly altered genes in LUAD, encodes a tumor suppressing protein that maintains genomic stability by regulating DNA repair, cell-cycle arrest, and apoptosis.^42,43^ Kurihara *et al*.^44^ found *TP53* mutations were associated with poor RFS in patients with primary stage I-III LUAD and La fleur *et al*.^45^ found mutations in *TP53* were associated with worse overall survival or a trend for worse overall survival in LUAD patients. *KRAS* encodes a small GTPase transductor protein that regulates cell proliferation and differentiation.^46^ In a retrospective study of patients with LUAD treated between 2008 and 2022 at Stanford University Hospital, Jiang *et al*.^47^ found patients with *KRAS* mutations had more aggressive tumors with a higher tumor mutational burden and had significantly shorter disease-free survival (DFS) and overall survival than patients with *EGFR* mutations. *DPP9* encodes dipeptidyl peptidase 9, a member of a family of atypical serine proteases that mediate a variety of processes, including cell migration, tumor cell invasion, and metastasis.^48,49^ Tang *et al*.^48^ found that high *DPP9* expression in NSCLC patients was significantly associated with lymph node metastasis and TNM stage and was a significant negative prognostic factor for 5-year overall survival. Additionally, repression of *DPP9* in NSCLC cells inhibited cell proliferation, migration, and invasion, while repression *in vivo* significantly reduced tumor volume and slowed tumor growth in *DPP9* knockdown nude mice. *VGF* encodes a neurosecretory polypeptide that plays a role in energy homeostasis and metabolism.^50^ In a study on EGFR-tyrosine kinase inhibitor (TKI) resistance in LUAD, Hwang *et al*.^51^ found that knockdown of *VGF* attenuated cell growth in EGFR-TKI-resistant LUAD cell lines and diminished tumor growth *in vivo*, while overexpression increased cell survival capacity, increasing migratory and invasive behaviors. Further, in an analysis of 70 LUAD tumor samples, the authors found a significant association between elevated *VGF* expression and poor survival. *NOTCH3* encodes a single-pass transmembrane protein belonging to the NOTCH receptor family, which plays a central role in the regulation of cellular proliferation, differentiation, and apoptosis.^52^In a study of patients with NSCLC treated with any surgical resection at National Taiwan University Hospital between 2001 and 2011, Chen *et al*.^53^ found that elevated *NOTCH3* expression in LUAD patients was an important predictor of poor DFS and associated with an increased risk of disease progression. A five-year prospective study of NSCLC patients by Ye *et al*.^52^ found similar results, with elevated NOTCH3 protein expression significantly associated with poorer overall survival in LUAD patients. Like *NOTCH3, NOTCH4* overexpression is a poor prognostic factor in a variety of cancer types.^54,55^ A recent multi-cohort study by Koh *et al*.^56^ reported that high *NOTCH4* expression was associated with worse overall survival in 1,161 LUAD patients from KMplotter. Authors found the same association between NOTCH4 protein expression and overall survival in a separate cohort of 347 LUAD patients who underwent surgical resection at Ajou University Hospital between 2009 and 2023.

While the results of this study are promising, there are multiple opportunities for future improvement and limitations to consider. First, after limiting to patients with recurrence data, both the primary dataset (TCGA) and external validation dataset (TRACERx) were small in size and thus prone to imbalance. As such, our findings were subject to a degree of noise. For example, even though LUAD is the predominant form of lung cancer found in never-smokers, the TCGA dataset had only 18 never-smokers (versus 114 past or current smokers), with only 1 never-smoker present in the TCGA validation set and 4 present in the TCGA test set. As a result, PLASMA struggled to learn accurate relationships between smoking status and patient outcomes. Second, to evaluate PLASMA’s generalizability to the TRACERx dataset, we were required to limit our clinical features to those available and populated in both datasets. As such, potentially relevant clinical features, such as surgical margin status, could not be used for training. Moreover, neither dataset contains comprehensive preoperative and postoperative treatment data, which would be expected to have a major modifying effect on the risk of recurrence. For example, perioperative chemotherapy plus immunotherapy strategies have been shown to improve event-free survival in registrational trials such as AEGEAN and CheckMate 77T and are now considered standard of care for early stage NSCLC.^57,58^ While the value of these public datasets as community resources cannot be overstated, these limitations underscore the importance of rigorously curated clinical data. Third, though we validated our model on the TRACERx cohort, there are notable differences between the two cohorts that may have limited validation performance. In particular, all TCGA patients were diagnosed between 1992 and 2013, predating the era of immune checkpoint inhibitors (ICIs), while TRACERx began recruitment in April 2014, just one year before the first FDA approval of an ICI for NSCLC.^20,32,59^ As a result, the two datasets reflect different treatment landscapes. Fourth, the TCGA dataset had more female patients than population averages (58% female) and both datasets had limited racial diversity, with 79.5% of TCGA patients and 95.5% of TRACERx patients self-identifying as white, suggesting selection bias. While PLASMA successfully generalized to the TRACERx dataset, additional external validation on datasets with greater demographic diversity will be essential to fully capture the model’s generalizability and reliability beyond the patients evaluated in this study.

At the modeling level, the PLASMA framework presents several limitations. First, the original genomic feature space was disproportionately large relative to the size of the training cohort, necessitating stringent feature selection to minimize the risk of model overfitting; however, such procedures can be overly sensitive and unstable when applied to small datasets. As a result, the model may inherit instability or biases introduced during feature selection, and its downstream performance and interpretability may reflect the inherent limitations of a reduced feature subset, including the potential exclusion of relevant signals and increased susceptibility to sampling variability. Finally, although PLASMA surpassed baseline comparators, there remains room to enhance its predictive power and performance before considering clinical translation. Despite these constraints, our findings demonstrate the feasibility of DL-based approaches to early cancer recurrence prediction.

In conclusion, this study highlights the capacity of DL to discern meaningful patterns within high-dimensional clinical and genomic data, even with constrained sample sizes. Additionally, PLASMA ablation results underscore the value of incorporating clinical, mRNA expression, and somatic mutation data for modeling recurrence. Future directions for this research include the incorporation of existing risk stratification models, graph-based approaches, and imaging data.

## Data Availability

All datasets used in this study are publicly available. The pre-processed datasets and all code are available at https://github.com/TranslationalBioinformaticsLab/PLASMA.

## Author Contribution Statement

Conceptualization: J.P.C. and E.D.U.; Data preprocessing, modeling, and data analysis: J.P.C. under the advisement of E.D.U.; Funding acquisition: E.D.U.; Methodology: J.P.C., C.G.A., R.S., J.L.W. and E.D.U.; Software: J.P.C.; Validation: J.P.C., R.S. and E.D.U.; Writing-Original Draft: J.P.C., K.H. and E.D.U.; Writing-review and editing: J.P.C., K.H., C.G.A., R.S., J.L.W. and E.D.U.

## Notes

### Competing Interest Statement

The authors have declared no competing interest.

### Funding

- Brown University Data Science Institute, Providence, RI

