## Supplemental Materials for "A Multimodal Neural Network Model for Early Recurrence Prediction in Lung Adenocarcinoma"

**Supplementary Methods**

**S1. Variant classifications within the TCGA and TRACERx mutation data**

During preparation of the mutation features, we filtered variant types to exclude silent mutations and RNA mutations. To do so, we utilized Variant_Classification in the TCGA annotation data and func and exonic.func in the TRACERx annotation data. Variant_Classification is a Genomic Data Commons (GDC) Mutation Annotation Format variable.^1–3^ Unlike TCGA, the mutation annotation data provided for TRACERx is ANNOVAR-formatted;^4,5^ however, the ANNOVAR variant classifications can be mapped to MAF variant classifications as shown in Table S1.^6–8^ To exclude silent and RNA mutations, we removed TCGA mutations for which Variant_Classification ∈ [“Silent”, “RNA”], and TRACERx mutations for which exonic.func ∈ [“synonymous SNV”, “synonymous”] (*silent*) or func ∈ [“ncRNA_exonic”, “ncRNA_intronic”, “ncRNA_splicing”, “ncRNA_UTR3”, “ncRNA_UTR5”] (*RNA*). The distribution of variant classifications in the TCGA and TRACERx mutation data prior to the feature-reduction and selection pipeline is shown in Table S1.

**Table S1.** Distribution of variant classification types in the TCGA and TRACERx mutation data.

|  | **TCGA** | | **TRACERx** | |
| --- | --- | --- | --- | --- |
| **Variant Category** | **Variant_Classification** | **N (%)** | **func / exonic.func** | **N (%)** |
| Missense | “Missense_Mutation” | 29,356 (76.75) | “nonsynonymous SNV”  “nonsynonymous” | 17,240 (43.48) |
| Nonsense | “Nonsense_Mutation” | 2,423 (6.33) | “stopgain SNV”  “immediate-stopgain” | 1,259 (3.18) |
| 3'UTR | “3’UTR” | 1,648 (4.31) | “UTR3” | 942 (2.38) |
| Splice Site | “Splice_Site” | 1,049 (2.74) | “splicing” | 542 (1.37) |
| 5'UTR | “5’UTR” | 951 (2.49) | “UTR5” | 919 (2.32) |
| Frameshift Deletion | “Frame_Shift_Del” | 932 (2.44) | Not present in TRACERx ☨ | - |
| Intronic | “Intron” | 800 (2.09) | “intronic” | 16,665 (42.03) |
| Splice Region | “Splice_Region” | 486 (1.27) | No equivalent | - |
| Frameshift Insertion | “Frame_Shift_Ins” | 239 (0.62) | “frameshift insertion” | 196 (0.49) |
| In-frame Deletion | “In_Frame_Del” | 97 (0.25) | Not present in TRACERx ☨ | - |
| 5'Flank | “5’Flank” | 94 (0.25) | “upstream”  “upstream;downstream” **§** | 275 (0.69) |
| 3'Flank | “3’Flank” | 80 (0.21) | “downstream” | 82 (0.21) |
| Translation Start Site | “Translation_Start_Site” | 48 (0.13) | Not present in TRACERx ☨ | - |
| Stop Loss | “Nonstop_Mutation” | 38 (0.10) | “stoploss SNV” | 24 (0.06) |
| In-frame Insertion | “In_Frame_Ins” | 8 (0.02) | “nonframeshift insertion” | 10 (0.03) |
| Intergenic | Not present in TCGA ☨ | - | “intergenic” | 604 (1.52) |
| In-frame InDel | No equivalent | - | “nonframeshift substitution” | 91 (0.23) |
| Frameshift InDel | No equivalent | - | “frameshift substitution” | 636 (1.60) |
| Unclassified | No equivalent | - | “UNKNOWN”  “unknown” | 166 (0.42) |

☨ Equivalent classifications do exist in the given format but were absent from the indicated dataset.

**§** When a resides within the downstream and upstream regions of two different genes, or within one gene for which the downstream and upstream regions (1-kb) overlap, ANNOVAR will classify the variant as “upstream;downstream”.^5^ There is one such case of this in the TRACERx mutation data, for a variant located in *H2AC20* (transcript size of 494 bp).

**S2. Correlation-based genomic feature reduction**

To reduce redundancy among molecular features prior to modeling, we implemented a two-stage, within-modality correlation filtering procedure. For continuous gene expression features, pairwise correlations were computed using the Pearson method, while binary mutation features were evaluated using the Jaccard similarity index. However, mutation features with fewer than 3 positive observations were excluded from analysis. Feature pairs exceeding a correlation threshold of 0.95 were flagged for removal. In the first stage, pairs involving at least one gene with prior literature support were resolved by retaining the known gene and discarding its correlated partner, provided the partner was not itself a known gene. The complete set of literature-derived LUAD-associated genes is detailed below.

In the second stage, remaining correlated pairs were resolved greedily by evaluating feature variance and, in the case of ties, correlation degree, defined as the number of above-threshold correlations involving that feature. Features with higher variance or lower correlation degree were preferentially retained as they were considered more informative and less redundant. This procedure was applied separately but jointly recorded across expression and mutation modalities, and all filtering decisions were determined exclusively on the TCGA training data.

**S3. Literature-derived LUAD genes utilized during correlation analyses and feature selection**

The complete literature-derived genes includes: *CCDC6*, *TP53*, *SLTM*, *PIK3CA*, *AQP2*, *CDPF1*, *LDHA*, *CD274*, *LMO7*, *TFG*, *SLC4A5*, *ATP1B1*, *PDGFA*, *CYP1A1*, *U2AF1*, *BCL2*, *NTRK3*, *PHACTR1*, *CCND3*, *GNPNAT1*, *GSTT1*, *DEPP1*, *SETD2*, *ALDOA*, *CDK4*, *FAM83A*, *STK11*, *SOX2*, *MDM2*, *MAP2K1*, *JAK1*, *MMP9*, *BRAF*, *NANOG*, *CAV2*, *HRAS*, *CD74*, *CHRNA3*, *TIMP2*, *RAD52*, *LRIG3*, *NOTCH4*, *TERT*, *TPR*, *NOTCH1*, *STRN*, *GATA5*, *CCNE1*, *EXO1*, *NRAS*, *MAP2K5*, *CHRNA5*, *CTNND2*, *SDC4*, *CDKN2A*, *PDGFRA*, *SMARCA4*, *TRIM33*, *ARID1B*, *DLGAP5*, *SLC3A2*, *TCN1*, *GATA4*, *KRAS*, *GSTP1*, *TRRAP*, *CLPTM1L*, *FOXRED2*, *POU4F2*, *ALK*, *RET*, *CLTC*, *GSTM1*, *HOXA9*, *KDR*, *KDELR2*, *ERBB3*, *MGAM*, *MYC*, *KEAP1*, *ASPM*, *CDKN2B*, *CD44*, *CDKN1B*, *DCTN1*, *FGFR1*, *ERBB2*, *MECOM*, *GCC2*, *SOX17*, *CYP2D6*, *KIF5B*, *ENTPD2*, *MMP2*, *NTRK2*, *EZR*, *SQSTM1*, *GATA2*, *CTNNB1*, *WIF1*, *ATM*, *FRRS1*, *POU5F1*, *HEY1*, *ROS1*, *BIRC6*, *RASSF1*, *NOTCH3*, *TPTE*, *GOPC*, *NF1*, *HYKK*, *VEGFA*, *NFE2L2*, *S100P*, *ABCG2*, *CNTNAP2*, *NOTCH2*, *CTLA4*, *RHOV*, *HIC1*, *PTPN11*, *STX2*, *EGFR*, *IGF1R*, *SPOCK2*, *RB1*, *PDGFB*, *EML4*, *CAT*, *ABCB1*, *HIF1A*, *NBPF1*, *ARID2*, *TPM3*, *RIT1*, *HEY2*, *NTRK1*, *PTPN3*, *CCDC174*, *SLC34A2*, *HOXD13*, *JAK2*, *TOP1*, *ZEB1*, *SFRP1*, *PTEN*, *TYMS*, *ABL2*, *ZKSCAN1*, *NRG1*, *CYP2E1*, *PCP4*, *COL1A1*, *NKX2-1*, *PROM1*, *FOXO3*, *PEX3*, *PDGFRB*, *ARID1A*, *GPX3*, *HIP1*, *HES1*, *ALDH1A1*, *CD109*, *KLC1*.^9–27^

**S4. Hyperparameter Tuning**

Hyperparameters were selected via stratified k-fold cross-validation (CV; k=3) on a development pool comprising the merged TCGA training and validation splits, with the held-out TCGA test set and the external TRACERx cohort left unseen during tuning. Within each fold, 15% of the training split was designated as an inner validation set to provide signals for early stopping and learning-rate scheduling.

The searches for PLASMA and its ablations used Tree-structured Parzen Estimator (TPE) sampling (applied via Optuna)^28,29^ over a discrete categorical search space comprising per-modality and fusion learning rates, scheduler factor and cooldown, and early-stopping grace period; encoder, fusion, and prediction-head architectures were held fixed across all variants. Trial budgets were scaled to search space size: 80 trials for the full multimodal model, 60 for two-modality ablations, and 40 for single-modality ablations. To minimize unnecessary computation, when TPE re-proposed previously evaluated configurations, identical parameter dicts were short-circuited and assigned the cached objective value. Among configurations tied within numerical precision of the top mean cross-fold AUROC, the configuration with the lowest cross-fold standard deviation was selected to favor stability across folds. The hyperparameter search spaces for PLASMA and its variants are available in Table S2, and the selected hyperparameters are available in Table S3.

Baseline classifiers were tuned using exhaustive grid search over their respective hyperparameter spaces, evaluated on the same stratified 3-fold CV protocol applied to PLASMA (including identical patient cohorts, fold assignments, and inner-validation split), so that comparisons across model families reflect differences in modeling approach rather than tuning protocol. Grid search was used in place of TPE because the baseline search spaces are small (24 to 72 combinations per model; Table S4) in comparison to PLASMA; exhaustive coverage is feasible within comparable compute and yields fully reproducible selection. Following PLASMA's selection protocol, configurations tied within numerical precision of the top mean CV AUROC were broken by lowest cross-fold standard deviation. Each model was subsequently constructed with its tuned hyperparameters, trained on the original training and validation splits, and evaluated on the held-out testing data (TCGA testing set and TRACERx).

**Table S2.** Hyperparameter search spaces for PLASMA and each PLASMA variant.

| **Hyperparameters** | **Search values** | **Applicable for PLASMA versions:** | | |
| --- | --- | --- | --- | --- |
|  |  | **Base/all-modality** | **Modality variant(s)** | **Simple concat.** |
| Clinical Encoder LR | {3e-4, 1e-4, 5e-4} | Yes | C, CE, CM | Yes |
| Expression Encoder LR | {3e-4, 1e-4, 5e-4} | Yes | E, CE, EM | Yes |
| Mutation Encoder LR | {3e-4, 1e-4, 5e-4} | Yes | M, CM, EM | Yes |
| Fusion/PredictionHead LR | {3e-4, 1e-4, 5e-4} | Yes | All | Yes |
| Early Stopping: Grace Epochs | {8, 10, 12} | Yes | All | Yes |
| LR Scheduler: Factor | {0.3, 0.5, 0.6} | Yes | All | Yes |
| LR Scheduler: Cooldown Epochs | {0, 1, 2, 3} | Yes | All | Yes |

LR = learning rate; C = clinical-only; E = expression-only; M = mutation-only; CE = clinical and expression; CM = clinical and mutation; EM = expression and mutation.

**Table S3.** Selected hyperparameters for PLASMA and each PLASMA variant.

| **PLASMA model** | **Clinical LR** | **Expression LR** | **Mutation LR** | **Fusion/PredH LR** | **ES grace** | **LR scheduler factor** | |
| --- | --- | --- | --- | --- | --- | --- | --- |
|  |  |  |  |  |  | **Factor** | **Cooldown** |
| Base/all-modality | 3e-4 | 5e-4 | 1e-4 | 3e-4 | 12 | 0.6 | 2 |
| Modality variant: C | 5e-4 | N/A | N/A | 5e-4 | 10 | 0.6 | 3 |
| Modality variant: E | N/A | 5e-4 | N/A | 1e-4 | 12 | 0.6 | 3 |
| Modality variant: M | N/A | N/A | 1e-4 | 3e-4 | 12 | 0.5 | 0 |
| Modality variant: CE | 5e-4 | 5e-4 | N/A | 3e-4 | 12 | 0.6 | 3 |
| Modality variant: CM | 1e-4 | N/A | 1e-4 | 5e-4 | 10 | 0.6 | 2 |
| Modality variant: EM | N/A | 5e-4 | 1e-4 | 5e-4 | 12 | 0.6 | 1 |
| No-attention/simple concat. | 3e-4 | 5e-4 | 1e-4 | 5e-4 | 12 | 0.6 | 1 |

LR = learning rate; PredH = PredictionHead; ES = early stopping; C = clinical-only; E = expression-only; M = mutation-only; CE = clinical and expression; CM = clinical and mutation; EM = expression and mutation.

**Table S4.** Hyperparameter search spaces and selected hyperparameters for the baseline classifiers.

| **Model** | **Hyperparameter search space** | | **Hyperparameters selected** | |
| --- | --- | --- | --- | --- |
| Logistic regression | C | {1e-2, 1e-1, 1.0, 10.0, 100.0} | C | 1e-2 |
|  | L1 ratio | {0, 1} (0 = L2 penalty, 1 = L1 penalty) | L1 ratio | 0 |
|  | tol | {1e-4, 1e-3, 1e-2} | tol | 1e-2 |
| Random forest | Max depth | {3, 5, 8, None} | Max depth | 5 |
|  | Min. leaf samples | {1, 3, 5} | Min. leaf samples | 1 |
|  | N estimators | {50, 100, 150} | N estimators | 100 |
|  | Max features | {‘sqrt’, ‘log2’} | Max features | ‘log2’ |
| Multi-layer perceptron | Alpha | {1e-5, 1e-4, 1e-3} | Alpha | 1e-3 |
|  | Learning rate init | {1e-4, 5e-4, 1e-3, 5e-3} | Learning rate init | 5e-3 |
|  | Hidden layer sizes | {(64), (128), (64, 32), (128, 64), (64, 32, 16), (128, 64, 32)} | Hidden layer sizes | (128, 64) |
| Support vector machine | C | {1.0, 10.0, 100.0} | C | 1.0 |
|  | Gamma | {‘scale’, ‘auto’, 1e-2, 1e-1} | Gamma | ‘auto’ |
|  | Shrinking | {True, False} | Shrinking | True |
| K-nearest neighbors | N neighbors | {10, 30, 50} | N neighbors | 50 |
|  | Weights | {‘uniform’, ‘distance’} | Weights | ‘distance’ |
|  | Minkowski p | {1, 2} | Minkowski p | 2 |
|  | Leaf size | {10, 30, 50} | Leaf size | 50 |

**S5. Model Ablation Statistical Analyses**

Statistical significance for each modality variant was assessed against the null hypothesis of chance-level discrimination (AUROC ≤ 0.5) using a one-sided label-permutation test. For each variant and test cohort, the null distribution of AUROC was constructed by randomly permuting the binary recurrence labels 1,000 times and recomputing AUROC against the model's calibrated ensemble probabilities; the one-sided p-value was the proportion of permuted AUROCs at least as large as the observed AUROC. The test statistic was the observed AUROC. Effect sizes are reported as the deviation from the chance baseline (AUROC−0.5), with positive values indicating better-than-chance discrimination. Two-sided 95% confidence intervals on AUROC were computed independently using a patient-level percentile bootstrap with 10,000 resamples; bootstrap iterations yielding single-class resamples (in which AUROC is undefined) were dropped, with the count of usable resamples confirmed to exceed 99% of the requested 10,000 in all reported analyses. Bootstrap CIs and permutation p-values used independent random seeds to ensure procedural independence.

Multiple comparisons within each cohort were addressed using Bonferroni correction. For the modality ablation analysis (Supplementary Table S5), p-values were corrected across the seven models tested in each cohort: clinical-only (C), expression-only (E), mutation-only (M), clinical and expression (CE), clinical and mutation (CM), expression and mutation (EM), and base/all-modality (CEM). For the fusion architecture comparison (Supplementary Table S6), p-values were corrected across the two variants tested in each cohort (CEM with GatedModalityFusion vs CEM-concat). Bonferroni was selected as the primary correction procedure to control the family-wise error rate at α = 0.05, consistent with the pre-specified small set of variants compared and the confirmatory nature of these comparisons. Statistical significance throughout this work refers to Bonferroni-corrected p < 0.05. Pairwise comparisons between variants were not formally tested because the size of the test cohorts (TCGA test *n* = 20; TRACERx *n* = 66) provides limited statistical power for paired tests to reliably resolve pairwise AUROC differences.

**Supplementary Table S5.** Modality ablation per-variant statistical analysis (one-sided permutation tests of AUROC vs chance)

| **Cohort** | **Variant** | **AUROC** | **95% CI** | **Effect size** | **p (raw)** | **p (Bonferroni)** |
| --- | --- | --- | --- | --- | --- | --- |
| **TCGA test**  **(n = 20)** | CEM | 0.850 | [0.643, 0.990] | 0.350 | 0.003 | 0.021 |
|  | C | 0.820 | [0.595, 0.980] | 0.320 | 0.005 | 0.035 |
|  | CE | 0.680 | [0.424, 0.901] | 0.180 | 0.100 | 0.700 |
|  | E | 0.630 | [0.364, 0.879] | 0.120 | 0.204 | 1.000 |
|  | M | 0.490 | [0.220, 0.766] | -0.010 | 0.542 | 1.000 |
|  | CM | 0.360 | [0.110, 0.637] | -0.140 | 0.853 | 1.000 |
|  | EM | 0.230 | [0.042, 0.467] | -0.270 | 0.976 | 1.000 |
| **TRACERx**  **(n = 66)** | CEM | 0.765 | [0.642, 0.874] | 0.265 | <0.001 | <0.001 |
|  | CM | 0.715 | [0.585, 0.832] | 0.215 | 0.001 | 0.007 |
|  | CE | 0.704 | [0.573, 0.822] | 0.204 | 0.004 | 0.028 |
|  | C | 0.697 | [0.565, 0.825] | 0.197 | 0.005 | 0.035 |
|  | EM | 0.636 | [0.491, 0.771] | 0.136 | 0.034 | 0.238 |
|  | E | 0.558 | [0.410, 0.706] | 0.058 | 0.216 | 1.000 |
|  | M | 0.461 | [0.321, 0.602] | -0.039 | 0.703 | 1.000 |

Test statistic: observed AUROC. Effect size: AUROC − 0.5. Degrees of freedom are undefined for nonparametric permutation tests; cohort sample sizes (n) are reported in lieu. Confidence intervals on AUROC computed from 10,000 patient-level bootstrap resamples. P-values corrected within each cohort using Bonferroni correction. C = clinical; E = expression; M = mutation; CE = clinical and expression; CM = clinical and mutation; EM = expression and mutation.

**Supplementary Table S6.** Fusion architecture ablation statistical analysis (one-sided permutation tests of AUROC vs chance)

| **Cohort** | **Variant** | **AUROC** | **95% CI** | **Effect size** | **p (raw)** | **p (Bonferroni)** |
| --- | --- | --- | --- | --- | --- | --- |
| **TCGA test**  **(n = 20)** | GatedModalityFusion | 0.850 | [0.643, 0.990] | 0.350 | 0.003 | 0.006 |
|  | Simple concatenation | 0.650 | [0.385, 0.893] | 0.150 | 0.147 | 0.294 |
| **TRACERx**  **(n = 66)** | GatedModalityFusion | 0.765 | [0.642, 0.874] | 0.265 | <0.001 | <0.001 |
|  | Simple concatenation | 0.533 | [0.387, 0.680] | 0.033 | 0.309 | 0.618 |

Same methodology as Supplementary Table S5. Bonferroni correction applied across two variants per cohort.

**Supplementary Figures**

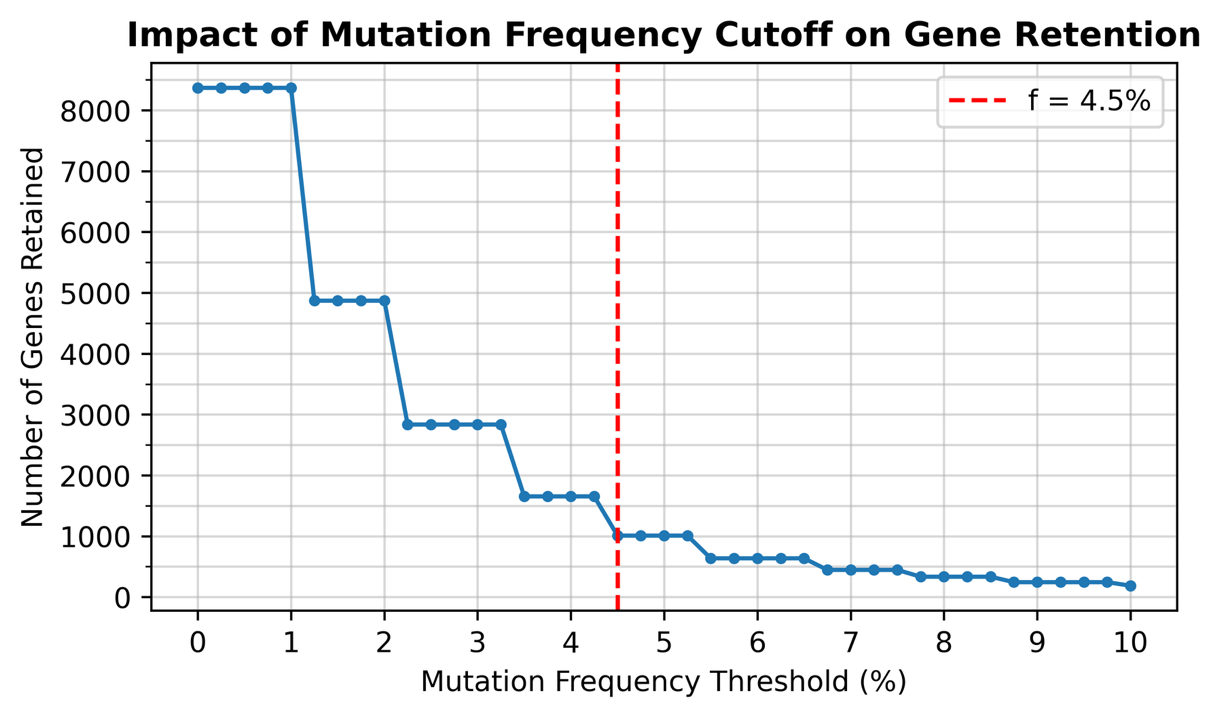

**Figure S1.** Impact of mutation frequency threshold on gene retention in the TCGA training set. Each point represents the number of genes retained when applying a minimum mutation frequency cutoff, calculated as the proportion of TCGA training samples harboring at least one somatic mutation in that gene. The dashed red line indicates the selected cutoff (*f* = 4.5%).

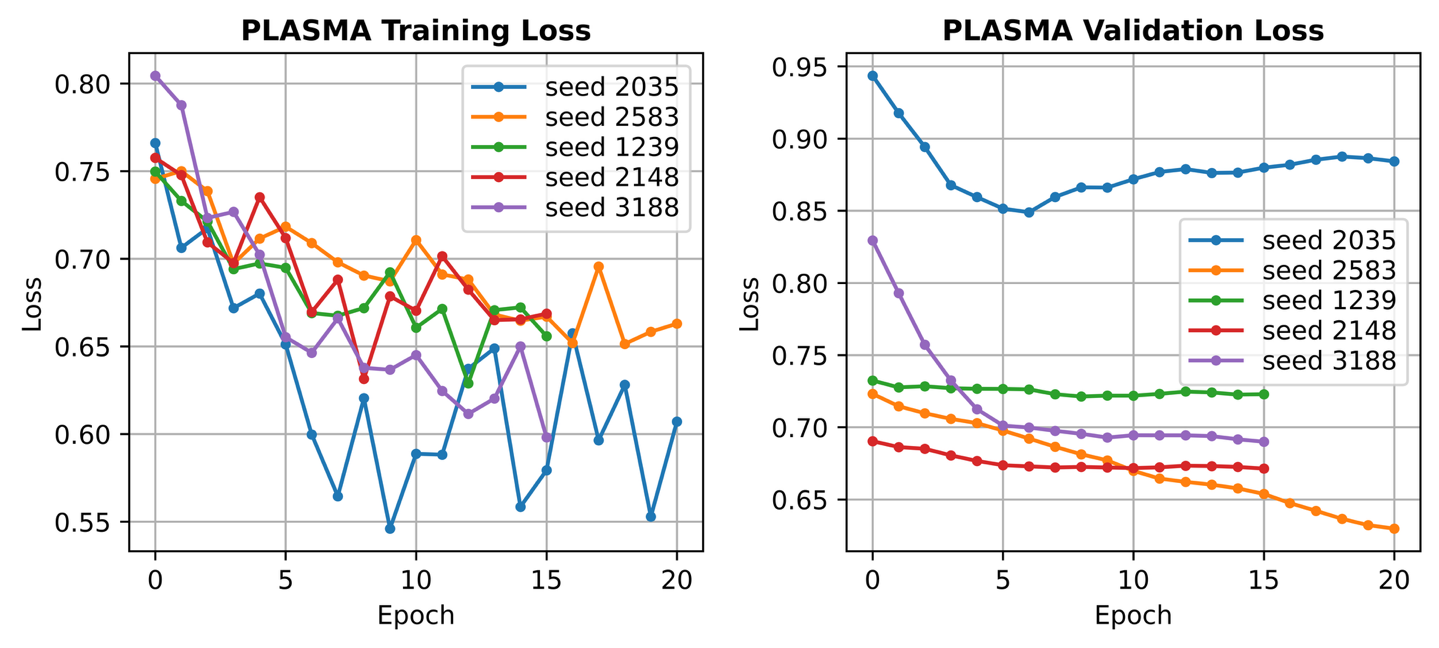

**Figure S2.** PLASMA training and validation loss curves across random initializations. Despite variability in initialization, all seeds show consistent downward trends in both training and validation loss, with only one seed showing evidence of overfitting. The spread in validation loss across seeds reflects sensitivity to random initialization rather than instability in the training procedure.

2.	GDC MAF Format v.1.0.0. *NCI Genomic Data Commons Documentation* https://docs.gdc.cancer.gov/Data/File_Formats/MAF_Format/.

3.	Mutation Annotation Format (MAF) - Legacy TCGA Specification (TCGAv2). *NCI Genomic Data Commons Documentation* https://docs.gdc.cancer.gov/Encyclopedia/pages/Mutation_Annotation_Format_TCGAv2/.

4.	Wang, K., Li, M. & Hakonarson, H. ANNOVAR: functional annotation of genetic variants from high-throughput sequencing data. *Nucleic Acids Res.* **38**, e164 (2010).

5.	Wang, K. Gene-based Annotation. *ANNOVAR Documentation* https://annovar.openbioinformatics.org/en/latest/user-guide/gene/.

6.	Mayakonda, A., Lin, D.-C., Assenov, Y., Plass, C. & Koeffler, H. P. Maftools: efficient and comprehensive analysis of somatic variants in cancer. *Genome Res.* **28**, 1747–1756 (2018).

7.	Mayakonda, A. maftools: Summarize, Analyze and Visualize MAF Files. (2018).

8.	Mayakonda, A. annovar2maf. (2023).
